## Supplemental Figures S1-S6 for "Regulatory genome annotation of 33 insect species": SupplementalLegends.docx

**Asma et al. 2024**

**Legends to Supplemental Figures**

**Figure S1: Lengths of predicted enhancers, including long outliers.** Size distribution of SCRMshaw predictions, prior to merging overlapping predictions but without removing outlier predictions. Species are ordered as in Fig. 2.

**Figure S2: Open chromatin data for predictions in the *ex, klu,* and *ush* loci.** SCRMshaw predictions (red bars) are shown for *D. melanogaster* (A, D, G), *T. castaneum* (B, E, H), and *A. mellifera* (C, F, I) at the *ex* (A-C), *klu* (D-F), and *ush* (G-I) loci. For *D. melanogaster* and *T. castaneum*, open chromatin profiles are also indicated, for a variety of tissues and timepoints (see text). Vertical gray bars highlight regions chosen for *in vivo* validation.

**Figure S3: Open chromatin data for predictions in the *hth, Ubx,* and *psq* loci.** SCRMshaw predictions (red bars) are shown for *D. melanogaster* (A-C), *Aedes aegypti* (D-F), *T. castaneum* (G-I), and *A. mellifera* (J, K) at the *hth* (A, D, G), *Ubx* (B, E, H, J), and *psq* (C, F, I, K) loci. For *D. melanogaster* and *T. castaneum*, open chromatin profiles are also indicated, for a variety of tissues and timepoints (see text). Blue bars indicate the positions of known enhancers. Vertical gray bars highlight regions chosen for *in vivo* validation.

**Figure S4: Additional expression observed in selected transgenic reporter lines.** (A-E) Control lines using a regulatory-inactive mock enhancer sequence demonstrate that there is no default expression in the imaginal discs. (A) Wing disc, (B) haltere disc, (C, D) leg discs, (E) eye-antennal disc. (F-I) Pupal expression. Arrows indicate expression in the legs (F-H) or notum (I). For *T. castaneum enhancer Tc_ex_9p0*, expression was observed in pupal legs but not larval leg imaginal discs (G, compare with Fig. 6B). (J) Expression can be observed in migrating adepithelial cells of the wing disc in *Tc_Ubx_19p9*. (K) Expression can be seen in the peripodial membrane surrounding the eye-antennal disc in several lines, including *Tc_ush_6p8* (see also Figs. 6, 7).

**Figure S5. Reporter gene expression observed in embryos.** A-S are putative enhancers cloned into piggyPhiGUGd and crossed to G-TRACE. T-Y have the putative enhancer inserted into piggyPhiGUGd-TomatoI. All embryos are stained using anti-dsRed antibodies and the ABC-HRP kit and shown with anterior to the left. (A-D) A non-enhancer control sequence was inserted into piggyPhiGUGd to serve as a control for vector-related expression. Limited segmentally-repeated expression is observed starting around stage 11 (A) and can be observed in the epidermis and/or peripheral nervous system at stage 14 (B). (C) Expression is observed in the caudal (longitudinal) visceral mesoderm (arrow, “cvm”). (D) In older embryos, strong expression is observed in the proventriculus (“pv”) as well as in migrating hemocytes (individual cells observed throughout the embryo). This “default” expression pattern was also observed in enhancer lines *Aa_hth_35p9* (E), *Aa_psq_21p5* (F), *Tc_Ubx_17p4* (G), and *Tc_Ubx_19p9* (H). Note that not all stages are shown for each of these genotypes, as expression was highly similar for each line at all stages. While this expression was seen in the remaining reporter lines too, additional activity, which may therefore represent specific enhancer activity, was also observed. (I) *Tc_psq_19p7* had prominent hindgut (“hg”) expression. (J) *Am_psq_29p2* had strong activity in the ventral nerve cord (“vnc”). (K) *Tc_hth_15p5* displayed epidermal expression following germ band retraction; note that the caudal visceral mesoderm expression can also be observed (L). (M-O) *Aa_Ubx_26p0* had reporter gene expression throughout the mesoderm (“me”) starting around stage 10 (M) and persisting throughout embryogenesis. “cb”, cardioblasts. (P) *Am_Ubx_0p39*, in addition to the default visceral muscle expression, was active in the ventral nerve cord (“vnc”) and salivary glands (“sg”). (Q-S) *Am_Ubx_37p2* showed expression in the epidermis from stage 11 on, and also in the salivary glands. (T-Y) Although we do not have a no-enhancer control line for piggyPhiGUGd-dTomatoI, all lines generated using this vector had reporter gene activity in the caudal visceral mesoderm (“cvm”), the salivary glands (“sg”), and in segmentally-repeated epidermal cells (arrowheads). We therefore view this expression as possible vector-specific expression (not all lines shown). However, two lines had additional, unique activity: *Am_ex_20p3* (X) was active in the ventral nerve cord, and *Am_klu_20p2* (Y) was active in narrow stripes in the epidermis (arrowheads). These activities may therefore be enhancer-specific.

**Figure S6. Revised post-processing method used for SCRMshaw.** See Methods for details.
