## Supplementary figures and images for "Regulatory genome annotation of 33 insect species"

### FigS1.pdf

Supplemental Figure 1

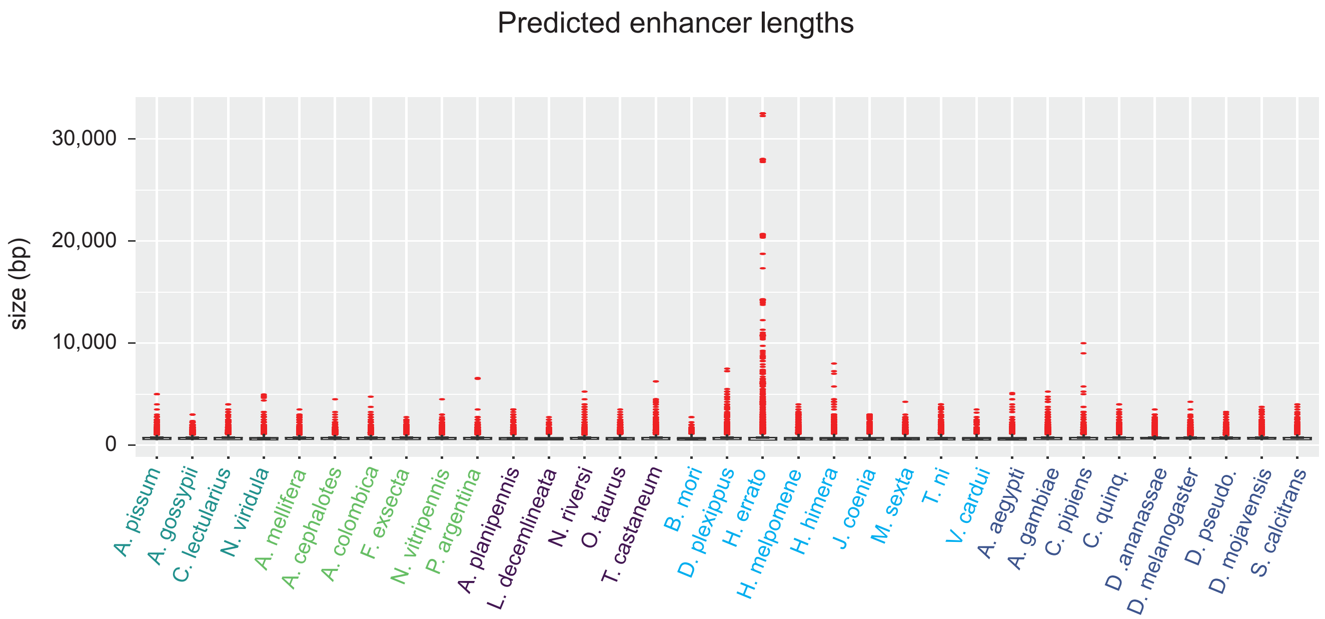

### FigS2.pdf

Supplemental Figure S2

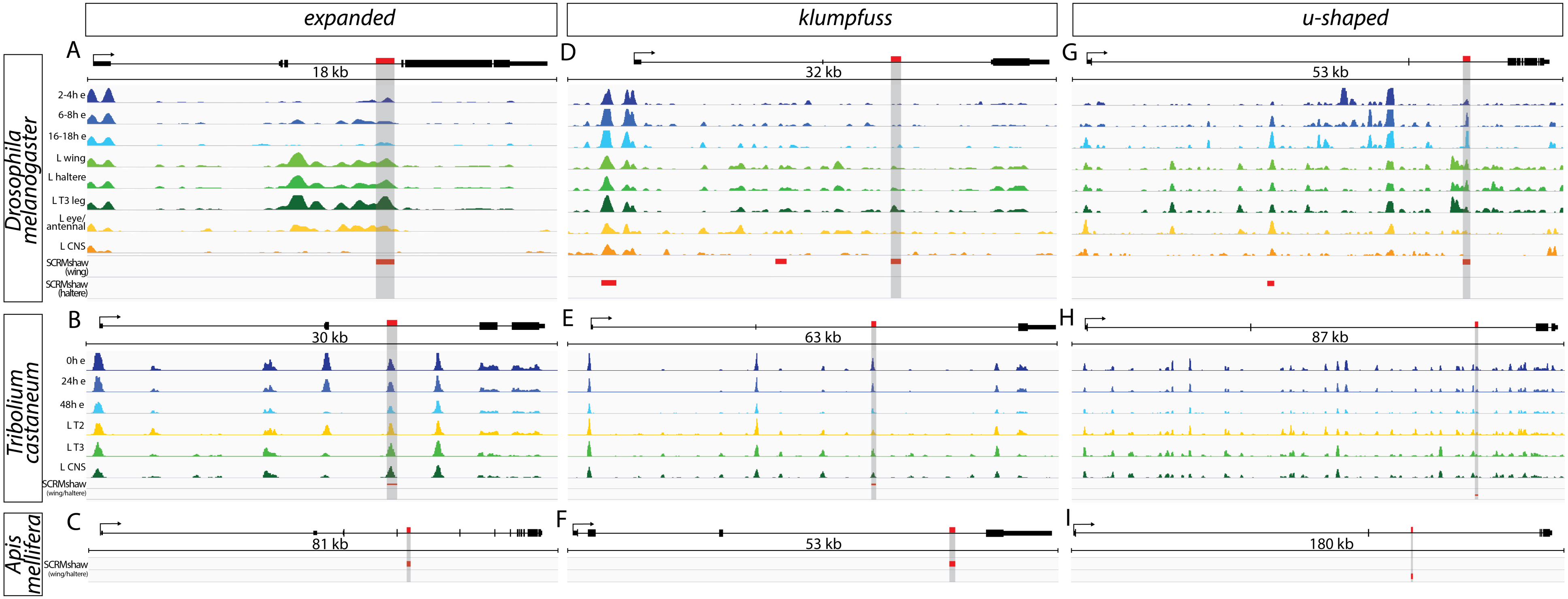

### FigS3.pdf

Supplemental Figure S3

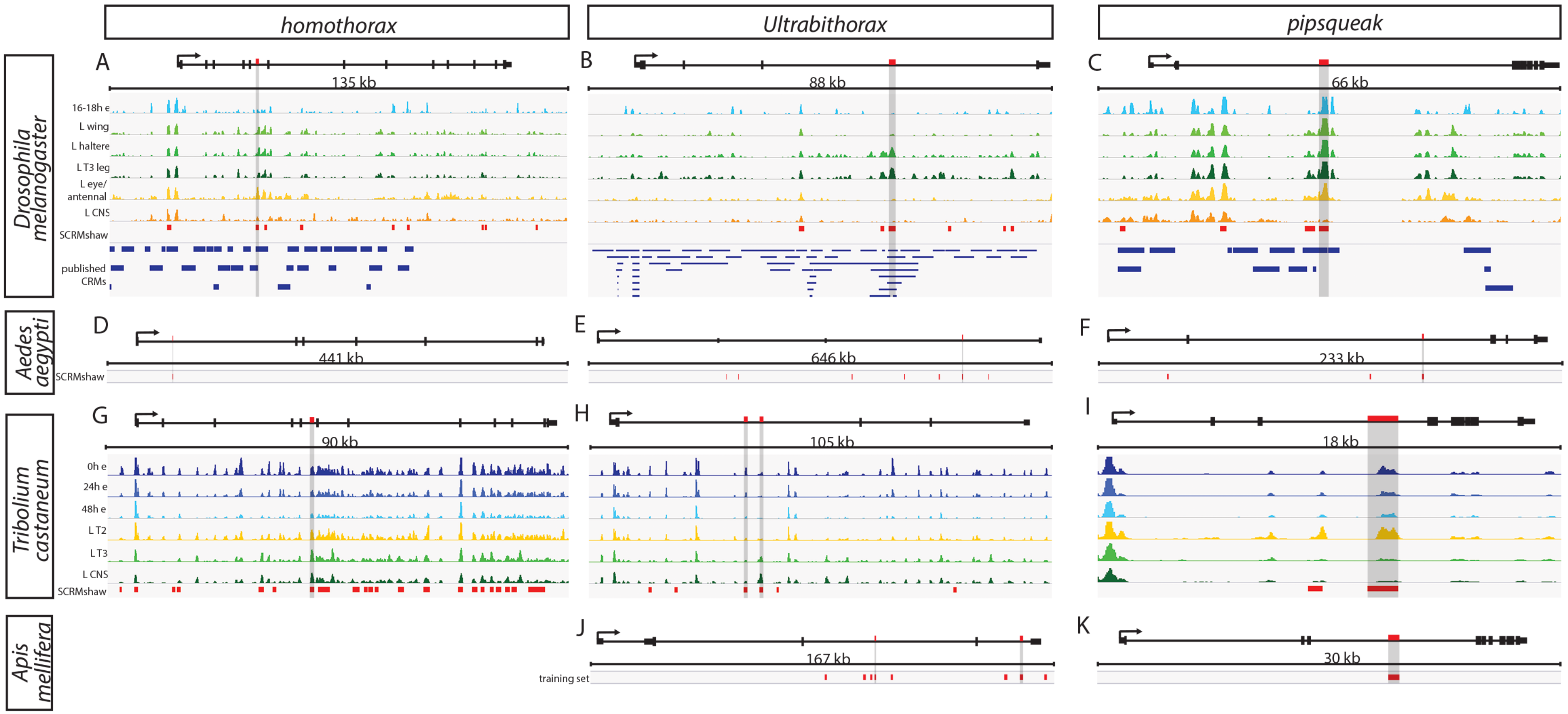

### FigS4.pdf

Asma et al. 2024 Supplemental Figure 4

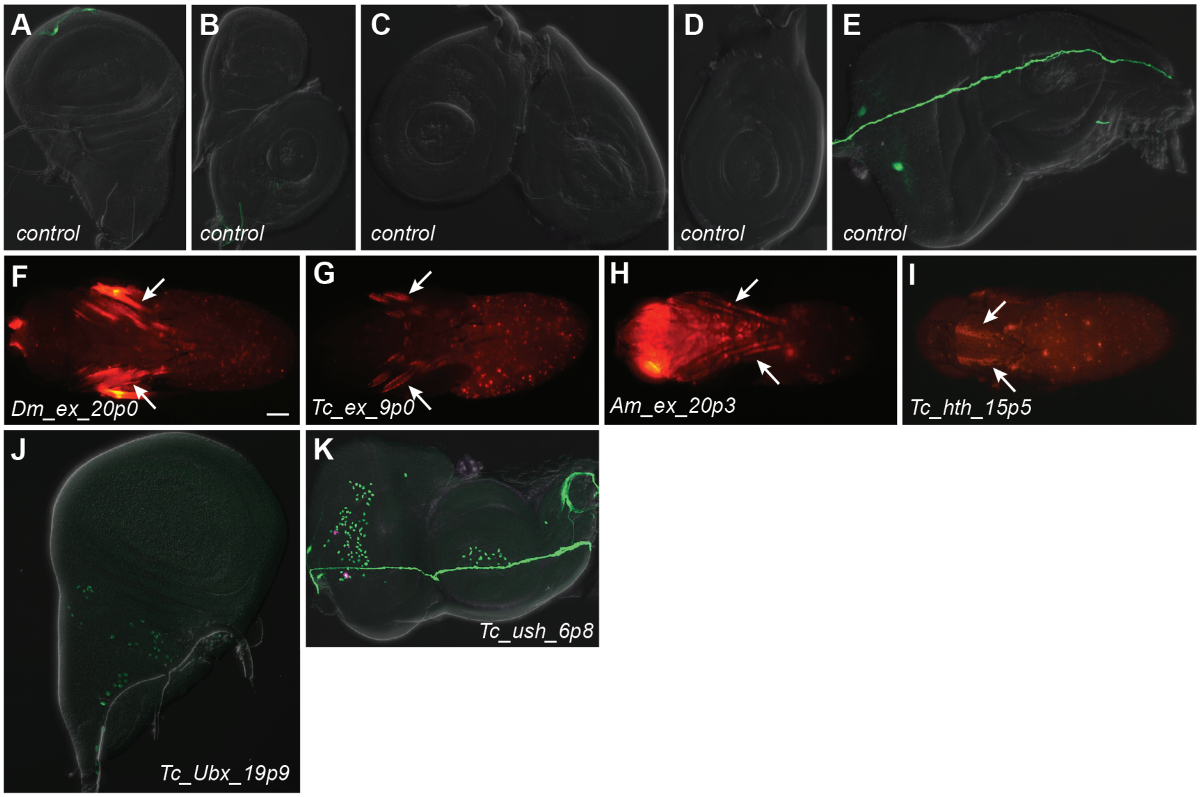

### FigS5-op.pdf

Asma et al. Supplemental Figure S5

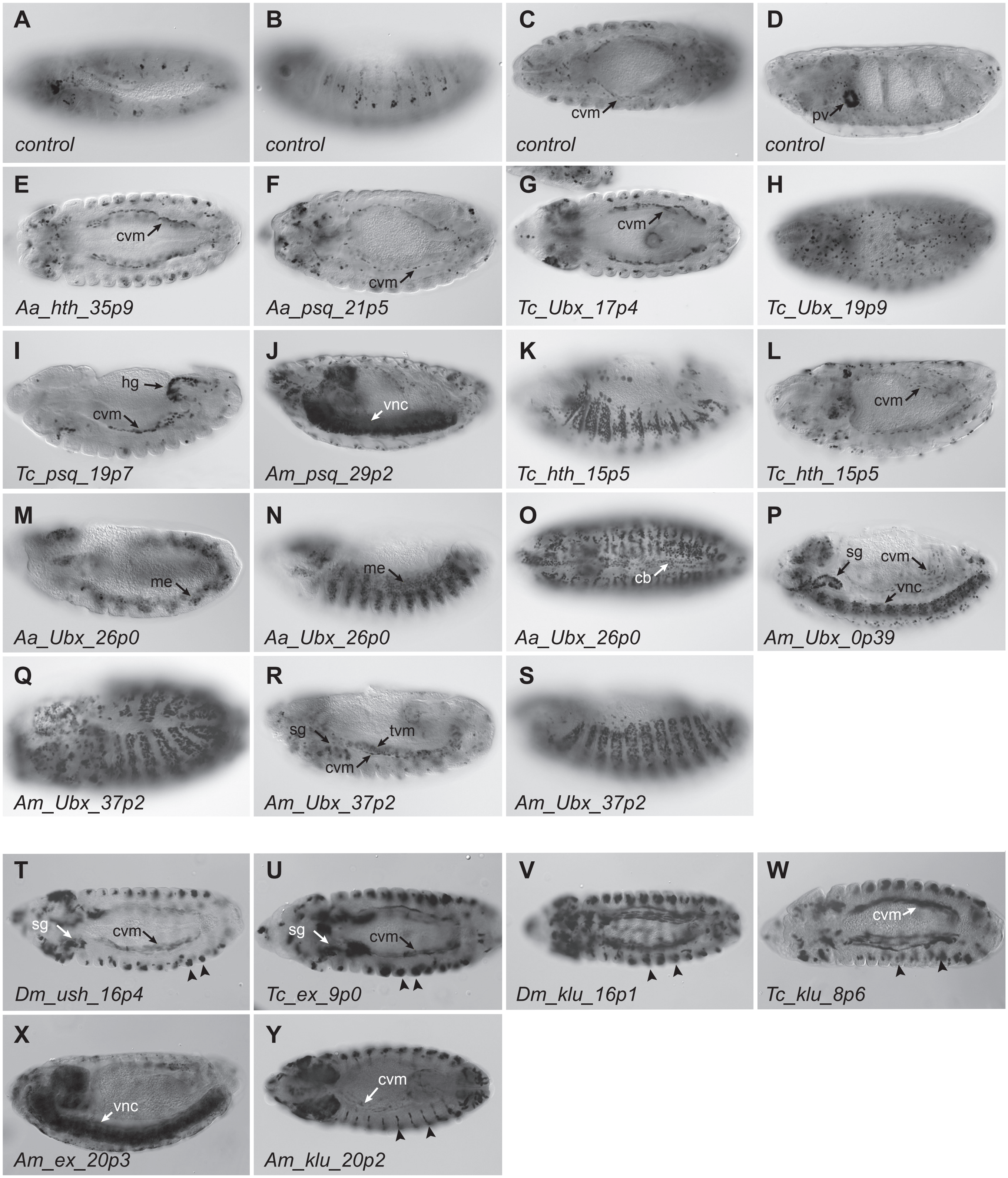

### FigS6.pdf

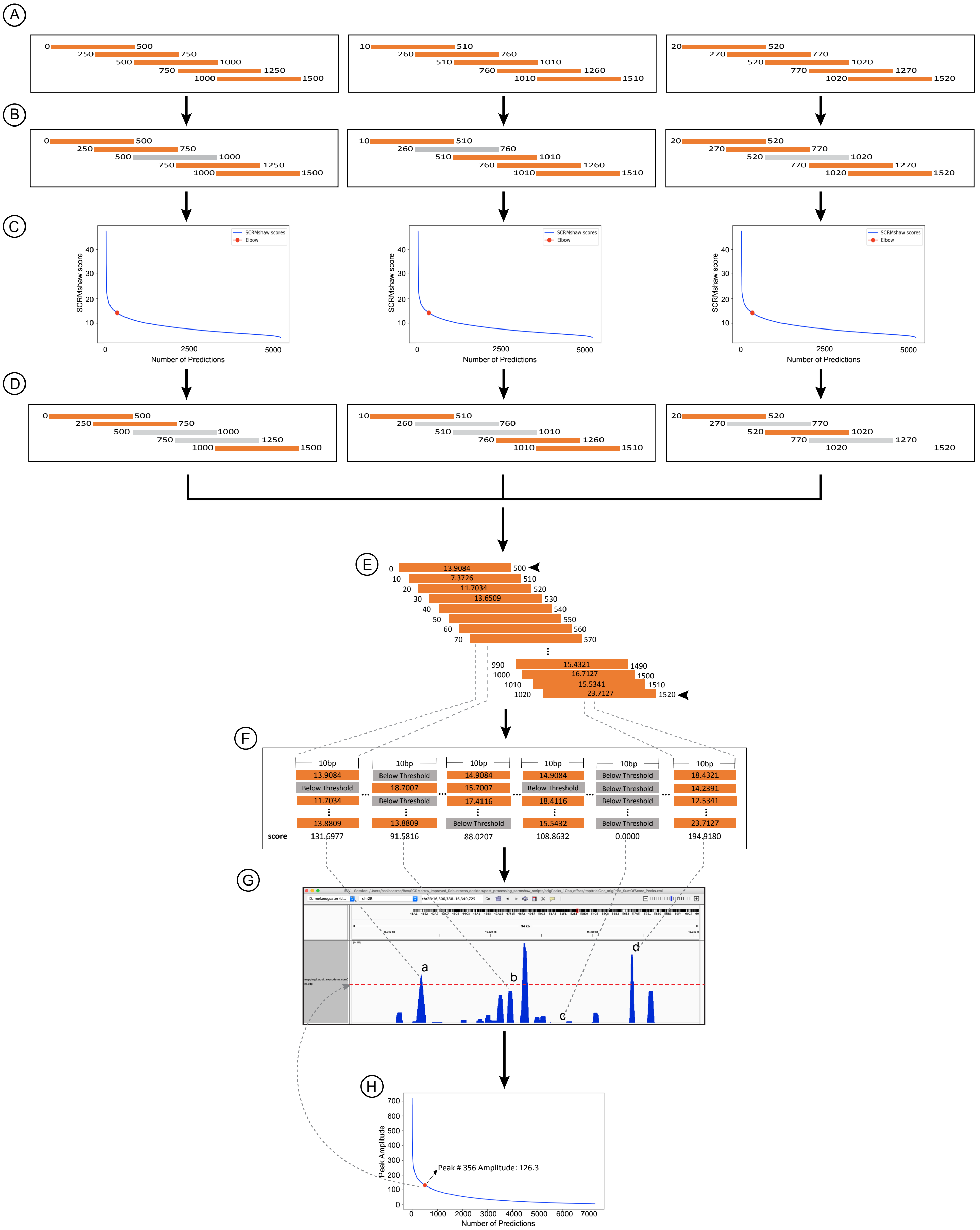
