## Supplemental Tables S1-S5 for "Regulatory genome annotation of 33 insect species": SupplementaryTableS5.docx

Table S5. Enhancer sequences predicted by SCRMshaw.

| **Fragment name** | **Size (bp)** | **sequence** |
| --- | --- | --- |
| *Dm_ex_20p0* | 700 | TTCCCAGAACAAACTTGTGGGGGGTGATTAGGTTTGGCAACAAAATATTTTGCTAGTATTCCCTAATCATTTTTTTGAGTGAACCAAACTCGAAGAGCTCTACTCCCCTGGCCATCCACTTGTTGCCACTTCCATTCCAGCTTTGCGTCGACGACGTCGTCATTGATAGGCACTTATTCGGCCGCTGATGATTATTATGATATTGTAGCTGCTGCTGCTGTGTTGTGGATTCGATGCTGAGGTGCCTCTATTCCATGGCCTCCTTCAACCTGCCTGCCTGCTTTTTTCATAATTATTATTTTTCATCTTGCTGCTCTTCATTTTGTATGCAGGAATTCCAATTTTTCGTTCGATGAAGTGTGTGTTGATTTCGCTGTTGTTTTTTCTCTGCCTTCTCGAGCACCGCCGACATGCCCTTGGGCCCTTCTGCTTGGCTCGGGTCGGAGCTATGTAGCGCGGTCCGGTACCGGTCTCGTCTTCGAGCATCAGGCAATGGGCCTCTGACAACCTGACGTGTCGTCATCATCATCGTCTTCATCTGCTGGAGTCTCTGACTCTTGTTGATGTCAATGGGTTGCTTGTTTATTGCCTGACAAACTGACAGAAGTCTGGTCGGGGTCTCGATCCGATTTGAGCCCGATTTGGGACGCAAGAGGAGCGCTCCCTCTTGCATAGCCGAAAGTTCATTTAAAATTTTGAT |
| *Dm_klu_16p1* | 660 | GACCAGGCTGTTGCAGTTTCGTGTTGAAACCAGTTTGAATATATTTATTTTTATTTCCTGCGTCCCCTTCCCAATTTCTGTGGCCCTTTTAGGCGCCTCAGTTAGTCGGCAACGATAAGGCGGCAATGGTTTAATTTAGCTGCACCAGCGGCAGCAGCAGATGACGACAGGATCGTTGGGCCGGTCTACGTGCAACAGAAGTTGCTGCGCCGGCAGAAGCAACGGCAGCGGCAGCAACAGCAACAGCAAAACAAATGTGTCTGTATATCGCAGCTAAATTGACTTTGATCACGCGATCCCGAATCCCCCCCCCCATTTGGTCCGAGTTATATGGCCGATTCCAGGTTGCAGGCTCCAGGTTCTCGGGCGCGGCCTTTTGTGGCACAAACGGAAGTATGCTAAGCAACTTGTTGCTGCCGCAAAGGCAAAGCAGCAAAAGCAGCTGAAGGTGTATATTGCAAAAATAATTACATTTGATTGTAAAAGGCCAGCGTCTCTAGGCTGGGGACTCGAGATCGGACCTCGGCCTGCCAGAGAAAAATGTGCAACATGATTGCAGTTTACAGCCCCAGCAGCAGCAGCAGCAGCTAAAGCAGCAACAACAACAGCAGCAGCAGCAGCAAAACCAACAGATAAAATGCATTTACAATTGAATTTGAA |
| *Dm_ush_16p4* | 810 | GAATGTTGCTGCGGTGGCATGTTGTTGCTCGTCGAAGTTCAGCCGATGTTGCTGCTGCTGCTGCAGCTGCTGTTGCTGCCATTCCCACTATCAACCGATGGTAATCGAAGGAAGCGACATTTATGCAAATGCCAGGTTGTTTAATAAACGCAAATTATGAGCCCGGCAGCAACATGTTGCAGCAACAGTCGATGGCAGATTAGCGACATTCATACTTGCACTTGGGTCAATTTAAATTTGTGCAACAGTGGCAGCACGGCACGGCAGCAACTCCTTGCCGCAGCAGCAGCCGCTCCAGCAGCACATGAGATTGTGGGAGCAACAGGCAGCATTATTGTGTTCGGCCAAGATCGCAATTGATCAGTGTGTTGGTGCTGGTGTTGGTGTTGCAGTTGCAGTTGCAGTTGCAGTTGCAGTCGCTGTTGCTGTTGCTGTTGCTACCCGACGACAACAGTTGCTGCTGTGCTGGTGCTAGTGCTTGTGCTTGTGCTTGTGCTTGTGTTGCTGCTGATCAAGCGATCAAGCACCGCAGCCAAAAACAATCGGCGCTGAGCGTGCTCACACGAAATTTTCAAGTACTGCGACAATTTCCATGCCCCCAGCCGCTGCCTTGTTATCAGCGCGCCATGCAACAGCAACAGCGACTGCAGCACAGGCAACAGCAGCAACACATCTCAACAGTTGCATCAATTGCTCAACATTGAACTCTGGGCATGGGCCCAGACGATCACCCCTCCTGGGGACCCCCTTCGGTCGCCCCTGCCCCAGTCCCTTGTATCATTTGCACGTTTTTTAATTAAGACATCAAAA |
| *Tc_ex_9p0* | 640 | ATAGTTCTAAAGTTCTAAACTATTTGCAAATGTAAACAACGACCGACATTTTCAACATGTTCGGGTGTACGTCGCTTTGAATATGGAAAATGTGATTTTGTAGAGAAATTTGGTGGCGTAGCCGTTGCCAGCTCCAGTTTCTTAAGGCAGGCATCTGGTAGCGCGCATCACAGCAGGCCGGGCCAGGTCAATAAAAAATCGAGTCAGCCGTGGGCTAAAAAACGCGACTAAAAAAACAATGCAGAGCCGCGGTTAAAGACAGGTAGCCGAGCTAAGCGGTAGGGGAGAGGGGGATCAGGATCATTTGTATACTCGGGGAATGTGCCCGACCCGGTACACTCGATGCCAAAACGAACACGCCGACTTATACAGATGCCTATCTGACAATACGGACACTTTTAAAAAACACTTTTTCGGTTTTTAAAAGTTAAAGCCAACAAGCGCTGTGTTTTACACAATTCTTCCAATTTCCATTCCCAAGTTGCAAAAGTGAAACGTCCCAAAATATTTTGTCGCGCAAATCAAAAGTATATTTTATTCATGAGGAGCTCGTGTAATTTTTATGTAAATTTTAATTATTGATAACAAGGGACCATTGTTTGACGACACTTTCTTCGGCATGCGGAGCTCCTTGTTTTGT |
| *Tc_klu_8p6* | 600 | TTATGAGTTTGGTTATCGTCGGAATCGGGCATTCTTCCTTTGTATCCGATTTTTAGGTAACAAGCTAGAAATTCCAGAGCTACACCTACGATCCATCAAGTCGGAGCCGTTCTAATTGGCCGCTCCTATCTATCGTCTGAAGGAGGCAGCAAGCAGCACACGAACACCTGGCTGCCAATGCACCTGATGCGGTCGTCCGTCGCTGCTATCAACTCACAATGTTTACTTGTGCTTACGCCAAAATTATGACGAATATTAATGCGGCCTCGTACGCAGGCAGGCATGCGGCGTAATTACTACCGGAGGGACCTTATCTCGAATTATTATGCAGCAAGCAGCAGTGAAAAATAGCGCAACGCCTGCTGCACTCATCCAGATCTGACAAAGATGAAGACGTCGCTGACATCTTTATGATTGTGCTTTTATTGCACCTTTTCGCGGAATTCCGACCATTCGAAGCGACCTTTGCGCTACGGAGAAGAGGAAATTTACACCGGGAGTTGACTTATGATGGGAGAACCACTCTCAACGAACGCAACTACTTTCCAGGAATATGAGAAGAGTGCTTAACTGACAAGTCCAAATTCGAACTTGAGGTTA |
| *Tc_ush_6p8* | 530 | TAAATAATCCAGACATGCATTGCATGTAAGTATCAGAGATACACGGTAAGAGTGGAGCTTTTCGAGAATCCGGAAACCGATCAGATAAGTTCTGAAAATGACTCGTCCACGAATAGATTTAGGATCGGAGCTGTTTTCCATTCGCCGAGATAAGTCGATAAGTTTCAATAAGTCCGAGTTCTGGCAACAGCCAGCACGGTACGGGCTGCCGCCGTCTTGGTTTCCAGTTTTCTCCAATGTCGTGGTATTAATCAGGGCGTTATCTCTAGCACATAAACACACGTATGTGTATGTGGGGCGATGTCGGTGGCGATACCGTTCCATGTGGGGGTGTAGCTGTTGGGGGTATACGGGCCTGTTCGCCGTCCGATAGCGCGAAAGATACGACCTGGAAGTAGGAAACGAGACAGCGAGATAAGAAAGTAATATGGCGGCTGCTGCAAAGAGATAACGACTGATACGCGCCTGCACCTTTCCCGACCTGCAACTCTACGTGCCCATTATTTTTGGAAAATTCAATGAGAAATCCG |
| *Am_ex_20p3* | 691 | TCTTGAGATCTTTCTGCATATAGCCGTGGTCTTCTTGCCCTCCCTCGCCCTCTGGCCCCGGACACCATCCACGGAGCTCCTTCCCTTCCCTTCACCGAATATACTCGGCTGTGCAGCGCCTGCTACCCTCTCGCTCTACTCTCTTGCCTTTCCACCAGTCATGACAAGCCGGTCCGACTGGTACCCCCACCAACGCGGCCGGACGGACCCTTCTTGGGCCTCGCAAGGGCCCTGCGGGACCCCTCCCTCCTACATTCCAGCGGGGCCCCATCACGGCGAGGCTGAGCTGGCGGGTTTTGAGGCGCGCGAGCCATGCCACGACAGGAAAAAAATGCATCTGAAAAACGAAAACAAGTAGAAAAAGGTGGCTCACACCCCTGCATGCGTGCGTCGGTTTGCGTGAACGTTGCCCGGACCCCGTACCGAGGCCTCCTCCTCCTCCTCCTTTCTCCTCCTTTCTTCCTTCCTTTCTTTTCGCTCCTCTACCTCCTCTGCGCGCCTCTTTACGCTCCTCTTCTTGCTCTACGCTCTCGCGTACGTGCCCGCAAACTGCTGCCTGCCTGTTCAACGCTTCTTCTTCGTTTCCTTCTTCTTCTTCTTCTTCTTCCTCCTCTCTGCTTCGTCCTTGCGTTTCTCTCGATTCGCGTCACATCTCCGCTCCCCCAAATACGTTTCCTTTCTAAGATCGTTT |
| *Am_klu_20p2* | 651 | TTATTTATCGCCCTCGAAAGCGCTCGTCCTCTGCAGATTTCGATCGAGTCGTTCGACTTCGATATAAGAATTTCAGTGTAAACGCGATACACGTTAAATAACGAATATTTACGGACAAAGTCGGGCGAACGACGCGATCCTGGCCGCTCGTGGCCGATGCGCAGGAAGGTAGGAGAGCGGAGAGGTTTTACGCTGTTCGCGGAGAGGAGGATAGGTTCGTATAGCTCCTTAAATCAACCCTAGTTGGCCTGTCAACCGAGTTGGCGCGCGCGCGCATTCCCTTTCGCGGCGCACAAATTACCACGCGTTTAATTACCGTCCGATATACGAAGCAGGCTCATTAATCACCACGCCGATAACCCGTAATTTTCCAGCAACGATAAAATCTATCGCGCGACACCGGCTCTCGCGACTTTCCTCTCTCTCTCTCTCCCTCTCTCTCTCTCGTTCGAGGAGAAAGGAGAAAAGGAAGCAGGAGGACGGAGGAGGTGTGCAGAGCGATCCTGTCGCCGCTTCCATATAGATTTTTTTTCTCCCTTCCGCGCTCTTTCTCGCGCCAGTTCTCTTCGTGCGGCGGAAAATAGAGCGGCGCAACTCCCCTTCTCGCGACTCACGGAGGGCGAACAGCTGAAGCCGGCCGATCGATACGAA |
| *Am_ush_20p8* | 651 | GGGGCTCCCTCCTCCTTCCCTCCTCCTCCGTCTGTCCCAGTTGGTCAGCCACGGTATCGTTTCGACGTCGATTCATCCCTCTTTTTCTCCCCCCTCTTCTCTCTCTCCTTCTGCCCCTCCCCCTCTCCCCTCCTCGCCATCGGCTTCGAGAGCCACGAGGCGATCGAGAGAGAGAGAGAGAGAGAGAGAAAGGGTACCCCATCGATGGATCGATCTATCGATCCACACGGGGATCCACCACGCTCTTCTGCCCTCTCCTCCTCCTCGCCACAATTTCTCTCCTCTTTACGTACGCTTCTCTCGTCCTTCGTGCCGCTCTTCGTCGCCATCGAGATTACGGCGAGCGAGGGGCCGACAGCCGAGGGGCTTCTTCCAATACTTTGTAAGTTTATTTGTATGATCCGCCAATACTTTGTATCTTTATTTATATGAAATCGGATGGCGGATCGAGATTGCTCTCTCTCTCTCTCTCGTGTCGCTCGTGTCTCGTCTCGCTTCTCCCCCCGTTTCCTCTTAAAATTAATTATACGTCCAAGGTGGGCGTAAGAGAGAGAGAGAGAGAGAAAGAAGTCGCAATGAAACCGGAAGGATAAAGAGAATCCGATGGTGCGCACACGCACGTGTATGTACACGTCCACTTTATAACACTCG |
| *Aa_hth_35p9* | 750 | TTCCGAACACCTTGATTCAAATCCGAACAGTAGGTACGAATAAATCATACCGTTTTGCTTCGAAATCTGGACACCTAAGACGAAGTGTATTTTCAAACTTGAATATATATATAGTAGAGATGGTCGGGTTTCACATTTTTCAAACCCGAACCCGACCCGTACCCGACTTATTTTATTTCTTCGAACCCGGACCCGACCCGAACCCGAGACCATAATTGAAAAGCAAACCCGGACCCGACCCGAACCCGAAAATTTTTCACAGTGCAAACCCGAACCCGACCCGAAACCCGAAAAATGTTTGTAAAAAAACCCGAATACAACCCGAGTTTGAAAAGATGGTAAATTCATCGTTTCTGATGCATAAAGAAGCTTTTAGATTGTTACTCTGTTCACAATTTTCACCAAACCCGACCTGAACCCGATTCAAACCCGACTTTTTGTAAGCCCGAACCCGACCCGTACCCGATAATTTCGTAGCCTACAAACCCGACCCGAACCCGAACCCGAAAAATTTCAAATATTCAAACCCGAACCCGACCCGAACCCGTCGGGTTCGGGTTCGGGTCGGGTTTCGCGTTTGAAAACCCGAGACCCGACCATCTCTAATATATAGGCAAATTCATACATACTAAAAATCCAATGGTGATTCTTGCTTCGAAATCCGGACAGCATGTGAGAGCCGATTCAAATATTGGACACATTTGCTTCGAATTCCGGACACTTCTATTTATCTAGTTTGTTCAAAGATTC |
| *Aa_ubx_26p0* | 647 | AAGTCGGGTTTAGTCGGGTTTGTGTCGGGTTTGGTGAGAATTGTGAACAGAGTGACAAACTAAAACCTTCCCTATGTATCAGAAACGATGAATTTATCATCTTTTAGAACTCGGGCTTTATTCGGGTTTTCCTTAGAAAACTTTTTCGGGTTTCGGGTCGGGTTTGGGTTTCAACAGCGAAAATTTTTCGGGTTCGGGTCGGGTCCGGGTTTGATTTTCAATTATTGTCTCGGGTTCGGGTCGGGTCCGGGTTTGAAGAAATAAAATAAGTCGGGTACGGGTCGGGTTCGGGTTTGAAAAATGTGAAACCCGACCATCTCTACTCTTCAGGTAGTCGAGAGTTGTTTTTTTTTATCTTTTATTTTTATTTTAAAGGCACTCTGTGCTCGTGCCCACTACTATGCCGAAATCAGTTCATCTGTATCTTCTTCACCGATTAAGATCTATTTTTAACTAATCTATATTTAAATCTACTTTCACTCTCTTCTACTCGTTTGCTCTCATACCGAGCAGGTAGGAGAGTGCTCTGCTGATAGTCCAATCGATTTCCATAAGCCATAGTTCCATTGCTCTTGCGGTGGTTCGTTTTGCCATGTTCCTGAGTCGTTTGAGGCTAGCTGCCTGCGAAGTGGGTCAGTTTGTCTCAG |
| *Aa_psq_21p5* | 890 | CCGAATTTGTGAGAGAGATAGGAGCCAATGTTTGAGTGATTCCCGCGAAGAATTGAAACCTATAAACGATTCCCACTAATTTTTGCAACATCTGTGATTTTTGATTTGATTTGAAACTGCAACTGACAGAAGATAATCAAAATACACTTTTTTCGCATTCGTACATCAATTGACAACCATCACTTGACACACCTGGCGATATGGACCAATAGGTCTGTGCCACAATAAGGGAGAGAAAAAAAAAAAAAAAGTGTGAAGCAAAAACACGCACATGTAACTTAAAGCACCACAAGAACCCTTTCAGCACCGGCCGCTTATGCTGATTTTATTAAAAAGCTTTATGCATACATGTACATAAGAGTGAGCATGCCGAAGCTCGAAAGTGTGTGTATGTGCGAATGCGCCAAAACACGATTATGTTTTCGTTTGTATTTCTTTCTTTTGCCGGCAAAAATTCTGTGTTTCGTTTTTTGATAGTAGGTAACTATGCCCACACAGTTACGGATCACACATAGTCATGGATCACTTTGGCGTTCAACATCGGATAACTCGCTCAAAACATATTTGCATGTGATGTAAACATATTTTTGCCAAGTCATAATATTTGTCTTCTGTCATTTGTAATATCAAATAATAAGCATAACATTAAACCGCAAAATAATGGTGTTTTTGAAAAATGTTTAGTTTGTATTGCCAAGCTATCATTAAATAGTCATTTATTGTAAGAAGTGCCATCAGCATTCTCCTATGCTTTTAGGTGATAAAATTCAAATATTATACATAATAGTTCCTCGTTCTCTTGAATTCAGTATGATTTCTTTGTTAGAAAACATTTTTCTTGTTTGTTGATACTGAATATTAGCAATTCCAACTAGTGATATTAGCAATTC |
| *Tc_hth_15p5* | 794 | TAATCTTTTAATTTAAAGCGTAGCTGAGCAGCTGGCTCTAATTCCACTTTCCTTATTTGGTTTCGTTGGTGTGGATTTTTGAAACGGATTATTTCGAGAAATAATAGTTTATTAGTGGTGGAAATAATGAATGGGTCTGGAGCGAGTTCCAGAGTGCGATTGGTTGGTTAGCGGGTAAATTTTTAAAAAGTGGGTGTCTTCTCCGACGGCAATTTAACGATCGTAACGACGTCGTCGCTAATTAGGCTCGTTGAGGCCGTCGCTAGATCGATAACACAGGCTGCGACATCGTCACAATGCACCGGTCGGGTTACACATCGGAGTCCGTCTCCCGGGGGCCCGTCTCAGATTCTCCGTATTAAAACACCGACATGTAAAAATATGGAAATTGCGCGCGGCAGAATGCGGTCCGATCAACCGGATGGCCATCGCGCATCGCTTTGCATTCGCAGCCGCATTTAAATTGCTAAAAGGGGACACTATCGAGCGGTCCATCTCTCTCGCAGCGTTGCGATATTATAATCTTGTTGCAAGGTAAATGCACATAACCGGTTACCCCAGACAGACGACGTCTTTGACACGAAAAAACCTGCCATCTATGTACAGCGGATCCTAATTTACGGCCTTATTCCATGTCATTAAGAGCATACGGGACGGACACGTTTTTAGGAACTTCGGACCCGACTTATCTCCGCGGACCGATAAGGAAATGTGCCTCTGGACACCTAACTTTGCCGACCAACAAAATCATAACGCTCGCTCTATGCCCATTGGGCAACACGAAAAAACCTGCC |
| *Tc_Ubx_17p4* | 870 | TGCATGTATGTCGAGTGGGTCCGGATGATGCGAACTCCCGCCGATTTCTTCGCAATCTGCAAATTCGCTCAAGTAGCTTAATAACAATGACAAAAGTGAGGCGGTATATTTCCGGCCGTCCGTTGAAAATTGTAATGATGTTATTAAAATTATGACGTGGCCGTGATGGTCGCCGAATTCTGGCGAAACGGCCGCGTAAAAACGGCACATAATTGGCTGACATTAAGATGTATCTGGAGATGTTTTTCGAATGCCTTCGTCCGGCGCGAATGCCTGAATAAGCGGCAAAGCTCGGAAAGCTCTTATAAATAAAAATGTACGGAGCCAATCAGATCGGCGAGTAAAAAGTACGTCTTTTCTTACACCAGAGGATCGCAGCTGCCGCAGAATCCGGTCGCGGATAAGAAATAAGAAGTGCTGCATAAATGCATTGATCATTCGCCGGGTCTCCGTCTGCTGTTCCTCAGCGAGAAAACGGGTTTAAGTCTGGATACTTTTGGCTCTCTGGAAAGTGCTTTTTGCATTAAGCTGCCGAGAGAGAATAAAGACGTTTGCGGTGTCGGACGGTGACCAATGCTGCTGCTGCTGCTCTGCCTTCCAAGTGCGTGCTTTAAATCTTCCACTTTGCAAGTAAATCGAGACGAACGCTGAATATTTTACACGAACACTGTTTATAGCCCAAATAACAGCCTTCCAAGGGCGGCCACCATCAAAAAATGGAGCGCTCAAACCCGAAATATGGGCGGGCGAAAATTATTCAAACCACAAAGCGAGGAAATCAGAAATTCAAAAATTGACGGCTTTCAACTCAGGACTGAATTTTTATAAATTTTTGTTCGCTACTGCAATTTGGGACAGAAAATTACATCT |
| *Tc_Ubx_19p9* | 780 | CGTATTTAAATATCGTTAGGTTCGATGGTAAAATTGGAGAAAATTGTCGCGCGCGTTTAAGACAAAGAAAATTCCCGTCGGGTTATCAATCTTGGGTTATCTGTACCCTCGGGCCGAAAAACTCTGTAAAGAAGAGACAAAAGGACGTGACAGTCCAATTTCCATTTCAGATCGAAATTGTTCGCCCCCCGGAAGTTTATCGGGGCCCGTTGGCGGAATTAATAAATTGGTGCGCGACTTAATTGCGGCGATAAAGAAGAGAAGAACACGAATGAGGGACGGCGACAAAAATATTATTTGCTCGTGAACGAGGAGGCAAAGGGCATTGATATCTCGTGCAACGCCGGATATTGGCTGCTTCTGGTCGCGGTTTGCGGGGCTTCTAAGACTGTGCAGGGTTTGGGGAGCGGCCCCGAGCTCGAGAGAAATTATGTACGAGGCATTGGGAGCAATATATCTCCGGCGGGACGTGCCAGACAGAGTAGACGGGGTATTATATAGGAAGGAAGGAACCTGAGGCCGGGGCCGGAGCCTCCTCGTCCCCAGGCGCTCGTCCCCCAGAATGAGACACTTGCCGCCAAGTCCACCGCCTTAAATTGTCATCTGAAGAAAGAAACTTCATTACGAACTACGCCCTCATTTCTTTGCGAGGCGATCCATCGCGCAAAAGCAACGCACGCATTTTGCAACAACTATTCAACCACTAAAATTAAACGAATTTCAAACCTATTCCGGATTAATGATTTCCTCCTCGATTCAAGCTAATTGGGTGTTTCCTAG |
| *Tc_psq_19p7* | 1250 | GGATTTTTTAGATAGATCATCAAGTTAAAAGTGCTTCGAATATATGTCATCAAAAATAAGATCAACTGATGGCTTTCTTTGCTTTATTCCCAATCTACTGTTAGAAAATCAACAACAACTAAGTTTTCTGTAAAATATAGTTCTTTCGGTGGCAAGAATAATATTATAATCGGGTTTCTTCTGCCTTATATTCTGTTTTCTTTGCTCCTATGTTAGTGCAAGTGTGTAACTTGGCGAACTCTTTCGAATTATCAAGGAAGTGTGAGTTTTATGAGAAAACAGCTAAAGTCGCCCCTAATTTGTTGACTTATTTGCTTTCGTTGGTTCTCCCGTTCTTTGGAGTATGTCGTCCGGTTTTTCTTATGAGCCATAATTACAAATTTCCATTTTCGGTTTTCGGCTCGCGTTCGTTTTGGAAAAGAGCGAATGTGCGGCGCGTTCATTTTCAATTTTGCGCGACCGTCCGACATTTTCCAATTTTCCGTGCAAGGACGAGGAGCGAGTGCAAAAAATGGCAGTCCTTGTCTGCAAAAAGCCCCAATTAAAACCGAAGTTGTAGTAGTGCGTGCGCCGAGCATTTCTCTCGATCTATCACGGGGTAGCAGCATCCCTCCGTAGGCTCACTCTCTGGCCAGTCTTAGTTTGCGCTTTCCCCGGAATTCACTGAAGGTCGTCGAGGTCGCAAGTAAGTACACAGTGCATGTGCACTTGCATGCATGCTTGCACTTTCTGTGCCCCCGCGCGCCGCCGCCGCCGCCGCATTAGCGTCTCTGTTTTGGTCCTTATATCCATCCGCTGTTCCCTTCTTCTGTCTATCCTTCAACTTCCTTCGCCGCTCGCCAGCTCCGGGACGCCACTCCATCATAAAACTGCGACCGCAAAAGCGACACTCATTATCGATTGCTCCAAGACGAATTAAAAGCCGCAGCGCTCCCCAAAACCGGGTTATTTTTTTCGGAATTTTGCTGGCTTGGAGCGGACTCCCAGACGATCCCCGGACTAATCCGGAGGGTTGCCTGGCGAGCGGCATTCGGCTTTAGGCTCCGGGGCACGCATTGGGGGAAAGTGATGCGGTGTCTGGACCAATCAATACCGGGTTCAAGGACGGCTTCTCTTATATGTGTATGTGAGCTTCCTTTTCCCGCTCGTCAAAACGGGACAAGACGGGAATTAATTGCACGACAATTGGGACGCCGACTCCACAGATGGGGCGACAAAATGGACGCAACGAACTAAATCTATTCACTTTT |
| *Am_Ubx_0p39* | 500 | TCTCTCTCTCCTCGAGTGTAGCATATATCCATTCCACCATCGATCGAGGATTTCGATCCCCCTTGGACTCATGCTGCGATATTCGATCGTCCCTCCCCCCACTCCTCCGCGCTCTCATTCGATCCTTCTTTTCTTCCCTCCCCCCCAACCACTTTGATCCTTCTCTCTCTCTCTCTTCCTTCTTTTTACTTCTTCTTCTTCTTGCTGCTGCAACTACCCGCTGCCTCTAACCGCTAGCCGGACAAAACATTTCTTAATTGGGTTTCGTTCGGAAAGAACCGTCCGATTTCGTTTCGCAAGGGATCCAGCCTGCTGCTTCTGCCGGTTTTACCGCGTCTCTACGTGGCTTCTGTCGTTCCCTCCTCGTTCTGCTTGCCTTTCCTTCGAACGATTATTTATTTCGTCGTTCGAATTCCTTATTTTTCCATCCTGTTATCCCTTATTGTAATAAAGTAAAAATAATTGAATTTTCCTTCGAACGAGCGAAGTTTGTTCTAATC |
| *Am_Ubx_37p2* | 1000 | AGAGAGAGAGAGAGAGAGAAAGTCAGGCAGACGGAGACAGAGGAATGGGTTGGGTAAGGGGGATAGAGTAGGGGCGGGAGGGCGTTCCACGGCACCCTGCATGGGGTAGCTTGCAACCTCACGCGACACTAGAGCCATCTATATCCCCGGAGATTTATGAGTTCCTGGTGCAGCGGCTGCTCGCAGCAACTACACACCACGCAGTATCGGGTCCGGTGTTGGTGCTGCCCCCTGTCGCGACGGGCGTGCTGTTGCCCCGCGGGGGTTACGCGCAATTCGCGCTCCGTGCAACGTCGCCCTGATAAAAAACTCTTGCGACTCGATCTCAATCCCGATGCTTCTGCGAACCTTCCCTCCGTTCTCGCGCCGTCTCGTCGTGCGCCCGTCGCGGTCGTCTCTCTCTCTCTCTCGCTCTCCGGTGTTGGCGGGCTATCGGATCTTCTCTCTCTCTCTCGCTCACTTGGTTCGCTTCTTGCTTTCGATGCGACGCGACGACCAGCGATCTCACTTTCTCTCTCGCTCTCGCTTTCGAGCTCGCACTTGAAATATCGATCATCATTGTGTTTCCTACGCATTTGTAGACCGCAAACGCGAAATTATTATGGGCCTGTGCACGTTTGAATTTCTTATATTCTTTTTTTTCCATTATACGCTAGGTTAGCGTAGATATAATTCTGCTAAATATAGTGAAGATAATTCGAATTTAAATTAAATTGAAAATTTTCGTATTACATAATACTGTTTCGTTATTTATAACTGATTAGAATATTTATTGATACCAAATGAAATTTTTGGTAAACTCTCGAACATTGTTTCATTCTTCTATATCGTATTGGTGAAAAATTACATCTCGATTTTTTCTCACGAACTTATATCGCGGTAAAAGAACTGTGGACAACTGTGCAGCATCTCCTCGCTCGATGAAGTCATTTGAACGAGCATTCCTCGGCCGATCTCAGATACAATCTCCTTCAAACAAAGAGCTCCATTGCCGCGTGCA |
| *Am_psq_29p2* | 691 | TCGCACGAGTACATAACGCTACCTTTGTCGCGTCGAAGGTAGAGGCACGATTCTGTCCTTTCCCGTTCTCTCGCGAACCTTGCATCCGTCTTCGTCTCGCTGTGGCCAAACGCGTGCTAGGTCTTCGTCTTCCACATTCCGTCTCGTTCGTTTCCGCACAGACTATATTTCTGTTCTCGTTTAGCCGCGGAAAGTCTTGCTCGCTCCCACGGGAACCACTCGTCGATGCTCGTCGCTTAACCGTCAGAGGCGAGCGCGCATTTCTCTCAAACACCGCAGACTTGCCTCTCCGCCGATCCCCGTTCCCACCCCCGGTGCTCGATGCTCTCTGTCACCCCTCCACCAAACGGACTCCTACCGGCCCGCTCCCCTCGCTTTGCGCCGCTTTCCACCAACCGTCCTGCCACCCGCCGGTTTTCAACCCCTTTCCCCGCTCTCTCGGCGACTGGTCAGGTGCGCTCGCTCGCTCGCTCGCTCCACGCGTACGCTCAATCGCTCTCTGTCCACCGCCGAGCACGCATCCCCCGCGAGTCTCTTCCTCGTTGTACGCGCTCGAGCGCGGATTCAATCCGTCCTTGTTCGTCGCGTCGGCGAATTTCGCGGCGTCCTCCGCCGCCGCCGCCGCCGCCGCCACCTCTTCCTCCTCCTCCTCCGCCTCCTCCTCCTGCTGATACTCCTCTTCCTCCTCGGT |
